# Alcohol use reduces the efficacy of anti-PD1 immunotherapy by disrupting anti-tumor immunity

**DOI:** 10.1101/2025.07.14.664729

**Authors:** Kristen N. Gilley, Jason Garness, Ali Khan, Brandon Carpenter, Mitra Shabrang, Wolfgang Beckabir, Chad Pecot, Benjamin Vincent, Leon G Coleman

## Abstract

Immune checkpoint inhibition (ICI) has improved clinical outcomes for certain patients with cancer. However, only a minority of patients have durable responses with underlying causes of differential immune responses across individuals often being unknown. Lifestyle exposures impact immune function and may subsequently alter the response to ICI. Alcohol use is common among cancer patients with known detrimental effects on adaptive immune function. However, its impact on ICI efficacy remains unknown. To determine if alcohol impacts ICI therapies, we performed a retrospective assessment of outcomes for patients receiving anti-PD1 ICI across tumor types and employed preclinical mouse models of ICI for lung and bladder cancer. Alcohol use reduced ICI efficacy in human patients treated with anti-PD1 for lung and bladder cancer (HR ∼2.0) as well as in murine models ICI in lung (LN4K1) and bladder (MB49) cancer. Alcohol reduced tumoral T cell numbers, promoting less productive Th2 and Th17 CD4+ phenotypes intratumorally and regulatory phenotypes in the periphery. In both rodent and *ex-vivo* human T cells, alcohol disrupted T cell activation and effector functions. Thus, alcohol use negatively impacts ICI efficacy warranting alcohol cessation for this patient population.

## Introduction

Immune checkpoint inhibition (ICI) therapy has shown significant promise for cancer care by offering the potential for improved response rates and reduced side effect profiles across many advanced and metastatic cancers *(1,2)*. This has resulted in approval for use in melanoma, lung cancer, bladder cancer, renal cell carcinoma, Hodgkin’s lymphoma, head and neck squamous cell carcinoma, colorectal cancer, and increasingly more malignancies *(3)*. However, only a minority of patients have durable responses to ICI treatment, with average response less than 30% across most tumor types *(4)*. There have been extensive efforts across the field to define determinants of response to ICI and to predict the likelihood of response for individuals *(5-8)*. This has revealed that features of anti-tumor immunity are associated with response. This includes immune checkpoint expression levels, T cell tumor infiltration, interferon (IFN) induction, T cell receptor (TCR) repertoire clonality, and tumor mutational burden *(9-12)*. However, it is usually unknown what factors on the individual level cause variations in these features that contribute to differential responses to ICI. Any factor that disrupts anti-tumor immunity could potentially worsen outcomes (e.g., neo-antigen presentation or T cell activation, tumor infiltration, and killing) *(13)*. An individual’s germline genetics can influence anti-tumor immunity *(14, 15)*. However, environmental exposures can also powerfully influence on immune responses *(16, 17)*. For instance, modifiable lifestyle factors such as obesity, smoking and alcohol use all impact immune function *(18-20)*, though their impact on ICI is unknown. By defining how modifiable risk factors impact ICI response, recommendations can be established to improve the likelihood of durable response to ICI.

Alcohol consumption is common and is a known immune toxicant *(20, 21)*. A surprisingly high number of patients with cancer use and misuse alcohol, though the impact of alcohol on immunotherapy outcomes is unknown. The NIH All of Us study (>15,000 patients) found that 76.4% of cancer patients drink alcohol (vs 66% in the general population) with 38.4% engaging in hazardous drinking (Alcohol Use Disorders Identification Test-Consumption [AUDIT-C] score ≥3 for women or ≥4 for men) and 23.4% engaging in binge drinking (>4 drinks/occasion) surpassing national averages *(22)*. The National Health Interview Survey (>34,000 patients) likewise found that 35% of patients with a cancer history exceed moderate drinking and 21% report binge drinking *(23)*. Most of the known influences of alcohol on immune responses are in the setting of infection and support an impairment of response. Alcohol disrupts immune responses to infections such as HCV, HIV, and pneumonia, increasing morbidity and mortality *(24-26)*. Chronic alcohol use has also been reported to decrease the number of T cells and promote T cell apoptosis and reduce CD4+ Th1 polarization after infection which is needed for anti-tumor responses *(20, 21, 27, 28)*. Given the potentially detrimental effects of alcohol on immune responses and high rate of alcohol use in cancer patients, it is essential to understand the impact of alcohol use on ICI outcomes. As such, the American Society of Clinical Oncology (ASCO) released a statement calling for studies investigating the impact of alcohol on responses to cancer treatment *(29)*. We hypothesized that alcohol use would negatively impact the anti-tumor immunity to worsen ICI outcomes.

To determine if alcohol use influences ICI immunotherapy efficacy, we examined the impact of alcohol on ICI outcomes in humans retrospectively as well as in preclinical murine models. We found that patients with lung and bladder cancer reporting regular alcohol use had significantly reduced survival compared to from ICI onset. Mice that were treated with chronic alcohol showed reduced responses to ICI for lung and bladder cancer. Studies of immune cells *ex-vivo* and *in vitro* found that alcohol alters the tumor transcriptome and microenvironment, T cell phenotypes, and T cell function in a manner consistent with reduced anti-tumor immunity. The evidence presented here highlights a critical intersection between alcohol use and the efficacy of ICI therapy in cancer treatment. Given the prevalence of alcohol consumption among cancer patients, these findings have significant implications for clinical practice and patient education.

## Results

### Alcohol reduces ICI efficacy in human patients and mouse models

To test for associations of alcohol exposure with ICI response, we performed a retrospective cohort review of 238 patients receiving anti-PD1 ICI between 2014-2018 at UNC Chapel Hill Hospitals. Patients were subdivided into occasional (<1 drink/week) and regular alcohol drinkers (>1 drink/week) based on their self-reported alcohol consumption at the time of treatment initiation. Occasional drinkers consumed an average of 0.2 drinks per week while regular drinkers averaged ∼7-8 drinks/week (**Table S1**). No other demographic factors (age, sex, BMI, and smoking status) showed significant differences in all tumor types. However, there was a significant difference in drinking by sex in bladder cancer and smoking was increased among regular drinkers in head and neck cancer. Patients with lung or bladder cancer who drank alcohol regularly had markedly reduced overall survival from the time of diagnosis (HR: lung-1.63, bladder-2.56, **Fig. S1A-B**) as well as survival from the time of anti-PD1 ICI initiation (**Fig. 1A-B**). For lung cancer, occasional drinkers had a median survival of 4.0 years after ICI initiation, while regular drinkers had a median survival of 1.7 (HR: 1.70, **Fig. 1A**, log-rank \**p<*0.05). Multivariable analysis including age, sex, BMI, smoking and alcohol use found alcohol use to be a significantly and inversely associated with survival from diagnosis (β:-0.73, **Table S2**) and from ICI initiation (β: -0.75, **Table 1**). Alcohol use and smoking are often comorbid, and smoking was found to be positively associated with survival from ICI initiation (β: 1.4). When smoking was not included in the regression model the effect of alcohol use was slightly decreased suggesting an interaction between these variables. For bladder cancer, occasional drinkers showed a median survival of 1.5 years from ICI initiation, while regular drinkers had a median survival of 0.5 years (HR: 2.3, **Fig. 1A**, log-rank \**p<*0.05). Multivariable analysis found alcohol use was inversely associated with survival from ICI initiation when smoking was not included as a covariate suggesting a complex interaction between the two variables (**Table 1**). No impact of alcohol use on survival from ICI initiation was found in head and neck cancer or melanoma, though a trend toward reduced survival from diagnosis was found for melanoma (**Fig. S1C-D)**. This indicates regular alcohol use is associated with robust reductions in survival in patients with bladder or lung cancer receiving anti-PD1 ICI.

**Fig. 1.**
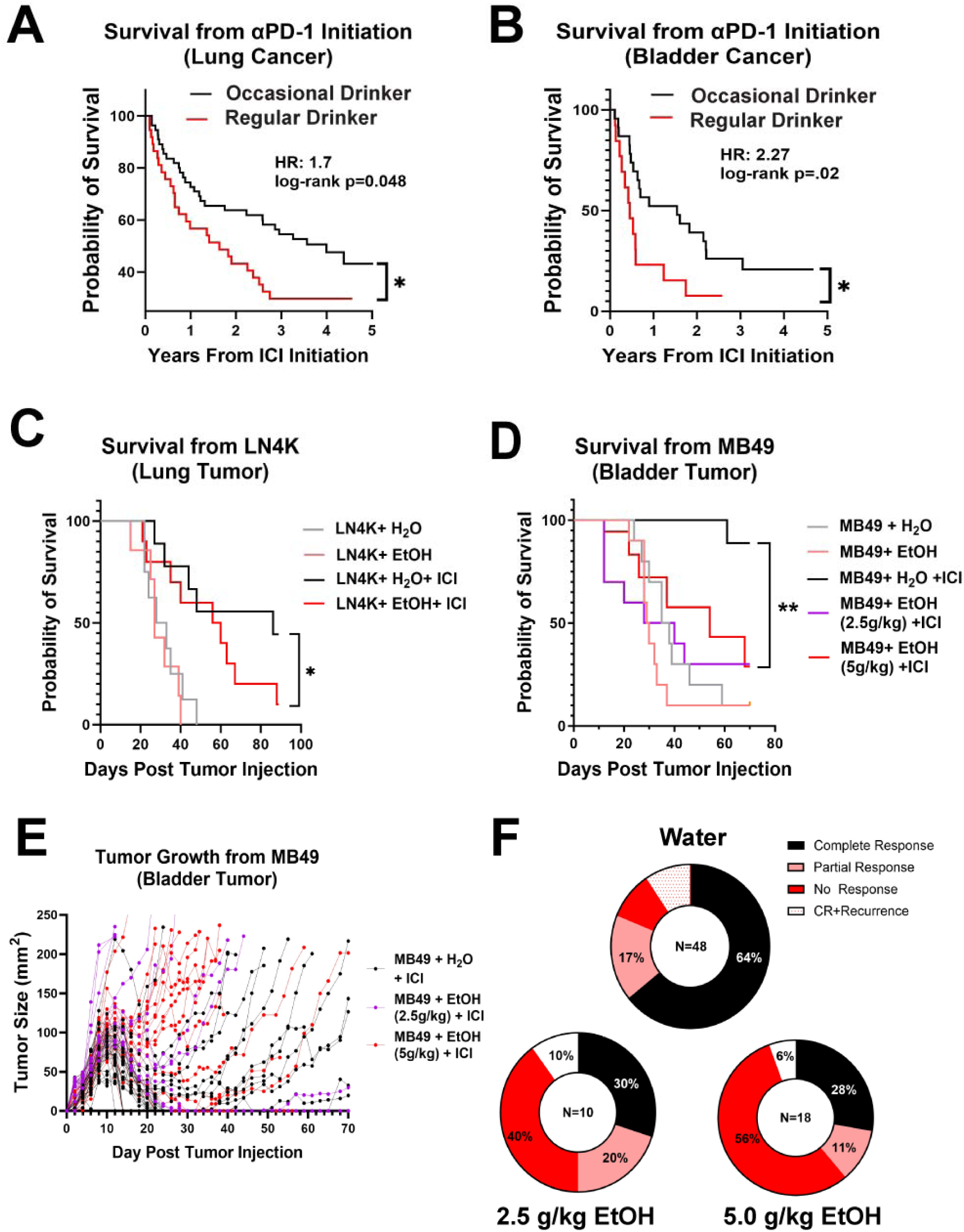
Alcohol decreases efficacy of anti-PD1 therapy in humans and murine models. (A-B) Retrospective analysis of human patients with lung (N = 91) or bladder cancer (N= 36) treated with ICI who were occasional drinkers (≤1 drink/week) or regular drinkers (>1 drink/week). **(A)** Patients with lung cancer who drank alcohol had worse survival from ICI onset than non-drinkers (HR:1.7, Log-rank test). **(B)** Patients with bladder cancer who drank alcohol had worse survival from ICI onset than non-drinkers (HR: 2.27, Log-rank test). **(C)** Mice received 5 weeks of water or ethanol pretreatment (5g/kg/d, i.g., 5 days on 2 days off) followed by LN4K lung tumor cell injection (10**^10^** cells/mouse, intrapleural). Water or ethanol was then continued until endpoint. Mice received ICI consisting of anti-PDL1 (10mg/kg) and anti-CTLA4 (10mg/kg) twice per week. Ethanol reduced the efficacy of ICI. **(D-F)** Mice received 5 weeks of water or ethanol pretreatment (5g/kg/d, i.g., 5 days on 2 days off) followed by MB49 bladder tumor cell injection (10^7^ cells/mouse, intradermal flank). Water or ethanol was then continued until endpoint. Mice received ICI consisting of anti-PD1 (10mg/kg) on post tumor injection days 8, 11, and 14. **(D)** Ethanol blocked the efficacy of ICI, reducing survival to control (no ICI) levels. **(E)** Tumor size trajectories for individual mice showing differential response profiles complete response (CR), partial response, no response, complete response with recurrence. **(F)** Ethanol increased the percentage of non-responders and decreased the percentage of complete responders

**Table 1.**
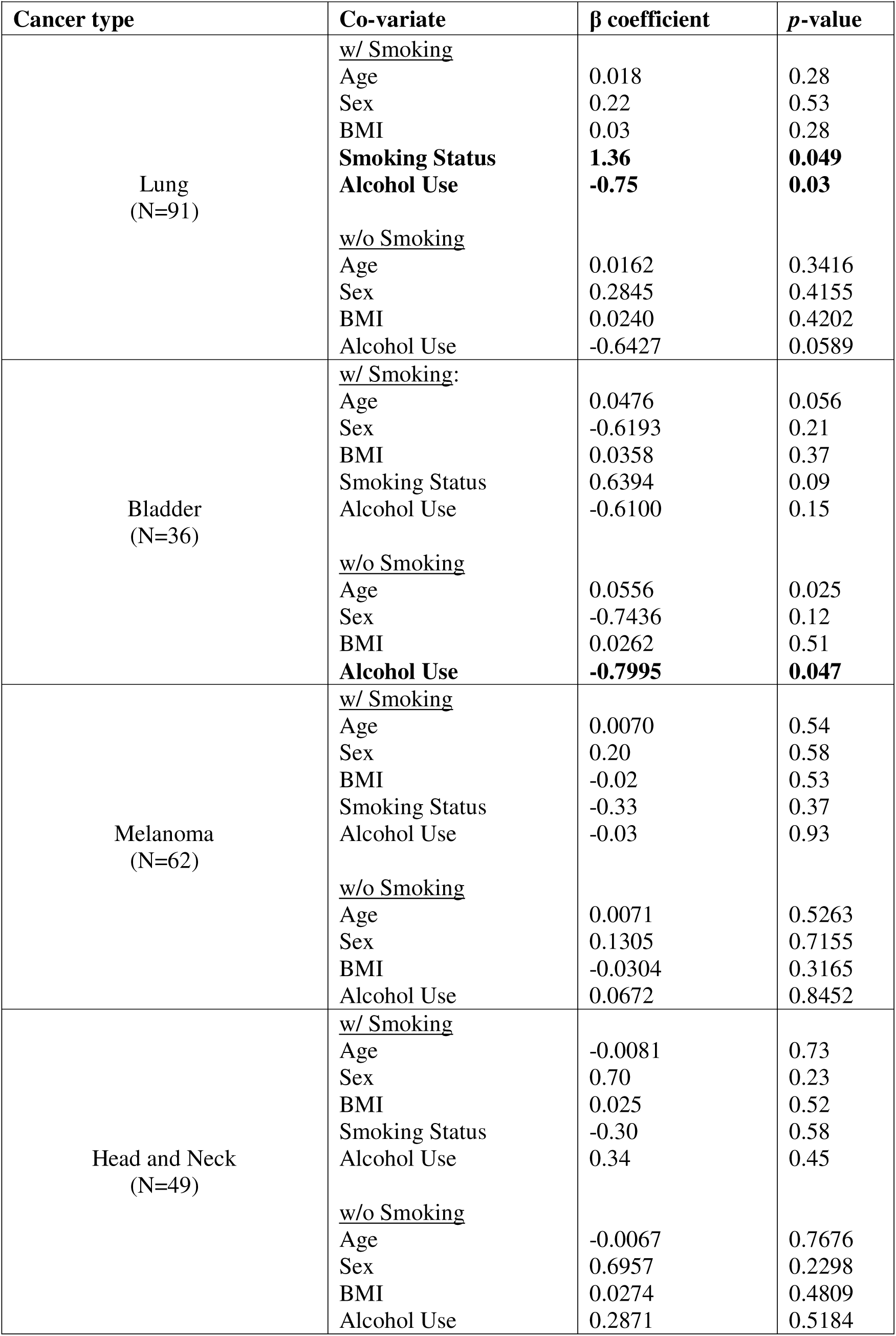
Impact of Alcohol Survival Post-Immunotherapy. Multivariable Linear Regression Analyses.

Retrospective analyses of human data can be limited by reporter bias or other occult factors. To determine if alcohol is responsible for reductions in ICI efficacy, we employed murine models of lung (LN4K1) and bladder (MB49) cancer. Mice received 5 weeks of alcohol pretreatment or water gavage in an intermittent pattern (5 days on, 2 days off) with continued treatment until endpoint to model human consumption patterns. Either moderate (2.5g/kg/day, i.g.) or heavy alcohol regimens (5g/kg/day, i.g.) were assessed. Mice metabolize ethanol up to 8 times faster than humans *(30)*. Pharmacokinetic studies comparing alcohol metabolism in mice and humans find 2.5g/kg (i.g.) is equivalent to ∼3 standard drinks per 70kg adult while 5g/kg equates to ∼6 standard drinks *(31)*. For a model of lung cancer, we utilized the orthotopic lung squamous LN4K1 model *(32, 33)*, which caused 100% mortality due to metastases within 50 days of injection. ICI with a combination of anti-PD-L1 and anti-CTLA-4 monoclonal antibodies (mAbs) prolonged median survival from 30.5 to 86 days in water treated controls. Ethanol (5g/kg) significantly reduced the efficacy of this ICI regimen, reducing median survival from 86 to 58 days (**Fig. 1C**, log-rank \**p<*0.05). In the absence of ICI, ethanol (5g/kg) had no impact on survival, suggesting a disruption of anti-tumor immunity rather than an enhancement of tumor growth. A similar effect was found with the MB49 bladder cancer model receiving anti-PD1 ICI. Both moderate and heavy ethanol blocked anti-PD1 mediated tumor clearance decreasing survival from 89% in water-treated controls to ∼30% with alcohol (**Fig. 1D**). Similar to the LN4K1 lung model, ethanol alone had no effect on MB49 tumor growth or mortality in the absence of ICI. Assessment of individual tumor response patterns found that alcohol reduced the proportion of complete responders and increased the proportion of non-responders (**Fig. 1E-F**). Thus, alcohol reduces ICI efficacy in mouse lung and bladder tumor models recapitulating findings in humans.

### Alcohol alters tumor transcriptomes, reducing immunogenicity and adaptive immune responses

To define the cellular and molecular impacts of alcohol that may contribute to reduced ICI efficacy with alcohol we analyzed RNA sequencing data on lung tumors from patients in Fig. 1 and MB49 cells treated *in vitro*. In patients with lung cancer who regularly consumed alcohol there were 1,327 significantly upregulated and 439 significantly downregulated genes compared to patients who occasionally consume alcohol (**Fig. 2A**). Many of the most significantly altered genes *(p<*0.001) are long noncoding RNAs (**Fig. 2B**). Ingenuity pathway analysis (IPA) predicted reductions in pathways associated with immune cell function including granzyme A, IL-7 signaling, TRAIL signaling, and innate and adaptive immune cell communication (**Fig. 2C**). Multiple pathways relevant to ICI response were also predicted to be increased including oxidative phosphorylation ***(34, 35)***, T cell receptor signaling, lipid antigen presentation by CD1, and regulation of IL2 (**Fig. 2D**). Upregulated pathways related to ICI response included oxidative phosphorylation, T cell receptor signaling, lipid antigen presentation by CD1, and regulation of IL2 (**Fig. 2D**). Thus, a history of regular alcohol drinking disrupts adaptive immune and metabolic pathways associated with ICI efficacy. These observed changes are likely due to a combination of influences on tumor, stroma, and immune cells within the tumor microenvironment.

**Fig. 2.**
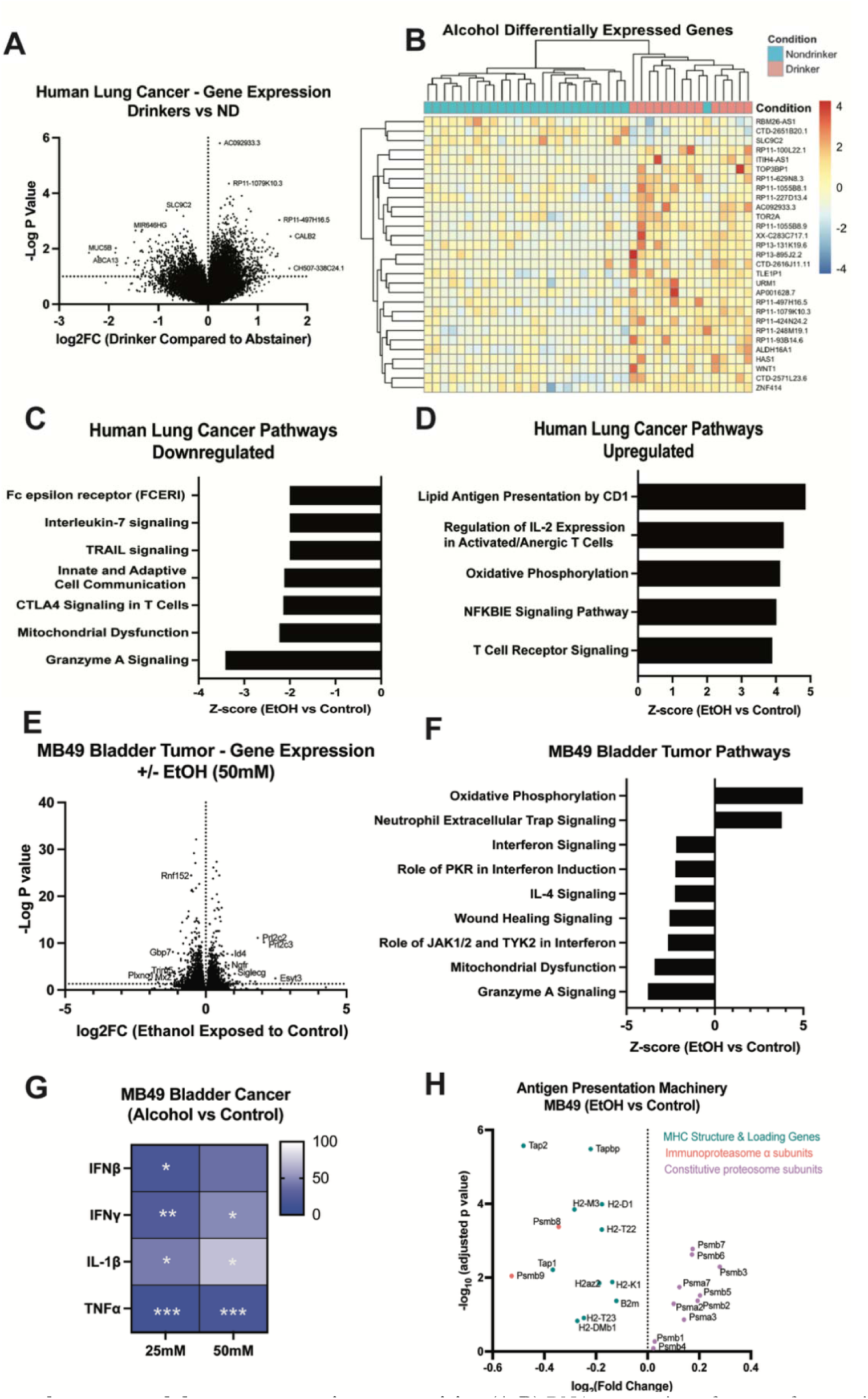
Ethanol alters tumor phenotype and decreases tumor immunogenicity. (A-B) RNA sequencing of tumors from patients with lung cancer, regular drinkers (>1 drink/week N=38) vs. occasional drinkers (≤1 drink/week, N=59). **(A)** Volcano plot of RNA sequencing of human lung tumors. **(B)** Heat map of most significantly differentially expressed genes of p < 0.001. (**C-D**) Ingenuity pathway analysis (IPA) of differential pathways (z-score greater than +/-2) identifying altered pathways such as increased oxidative phosphorylation and decreased Granzyme A, IL7, and TRAIL signaling. **(E-H)** RNA sequencing of MB49 tumor cell lines +/-ethanol (50mM, 24 hours). **(E)** Volcano plot of RNA sequencing showing several differentially expressed genes. **(F)** IPA depicts identifying altered pathways such as increased oxidative phosphorylation and decreased interferon response. **(G)** RT-PCR confirmed reduced immunogenicity found by RNAseq with reductions in expression of IFNβ, IFNγ, IL-1β, and TNFa (One-way Anova). **(H)** Volcano plot of genes from MB49 RNAseq associated with antigen presentation machinery finding a decrease in immunoproteasome subunit genes, while an increase in constitutive proteasome subunit genes.

To determine the direct effects of ethanol on tumor cells that showed reduced ICI response *in vivo*, we performed RNA sequencing on MB49 treated *in vitro* with ethanol. Ethanol had a robust effect on the MB49 transcriptome, significantly increasing 1,297 genes and reducing 1,286 genes **(Fig. 2E)**. Similar to human lung tumors, IPA predicted increased oxidative phosphorylation and decreased mitochondrial dysfunction **(Fig. 2F)**. Importantly, IFN signaling, which is essential for neoantigen presentation and induction of anti-tumor adaptive immune responses was reduced by alcohol. Reductions in IFNβ and IFNγ expression were confirmed by RT-PCR with concomitant decreases in pro-inflammatory genes IL-1β, and TNFα **(Fig. 2G)**. Since IFNs upregulate MHC and antigen presentation, we examined MHC structural components and immunoproteasome genes. Ethanol reduced MHC structural genes (B2M, Tap1, Tap2, H2 subunits) and immunoproteasome components (PSMB8 and PSMB9), while increasing constitutively active proteasome subunits **(Fig. 2H)**. Together, transcriptomic analysis of both human lung tumors and MB49 cells found that ethanol reduces tumor immunogenicity as evidenced by reduced IFN signaling and immunoproteasome component expression along with a predicted increase in tumoral oxidative phosphorylation, which can regulate antigen presentation *(34, 35)*.

### Alcohol reduces tumor T cell numbers while promoting Th17 and regulatory phenotypes

To define the impact of alcohol on immune cell phenotypes in the tumor microenvironment and periphery, we performed flow cytometry on tumors, tumor draining lymph nodes, and spleens of mice bearing MB49 tumors with anti-PD1 +/-alcohol. Mice were treated as in **Fig. 1D**, with sacrifice and assessment on day 11, the time of maximum tumor size just prior to either the beginning of tumor resolution or continued growth **(Fig. 1E**). Multiple cohorts were assessed, with consistency across experiments confirmed by a meta-analysis (**Fig. S3A-B**). Flow cytometry revealed disruptions in the adaptive anti-tumor immune responses. Within the tumor, there was a reduction in CD4+ and CD8+ T cells (**Fig. 3A-B**) with a significant increase in NK cells (**Fig. 3C)** but no changes in the number of B cells, dendritic cells, or macrophages (**Fig. S2A-C**). In the tumor draining lymph nodes, CD8+ T cells and CD4+ T cells showed trends toward significant reductions **(Fig. 3D-E**), and the proportion of B cells was increased (**Fig. 3F**) while dendritic cells, natural killer cells, and macrophages were mostly unaffected (**Fig. S2D-F**). To better understand the peripheral immune environment, we collected the spleen but found no significant differences in T cells, B cells, DCs, NK cells, or macrophages proportions (**Fig. S2G-L**). Thus, alcohol reduces CD4+ and CD8+ T cells within the tumor and tumor-draining lymph nodes.

**Fig. 3:**
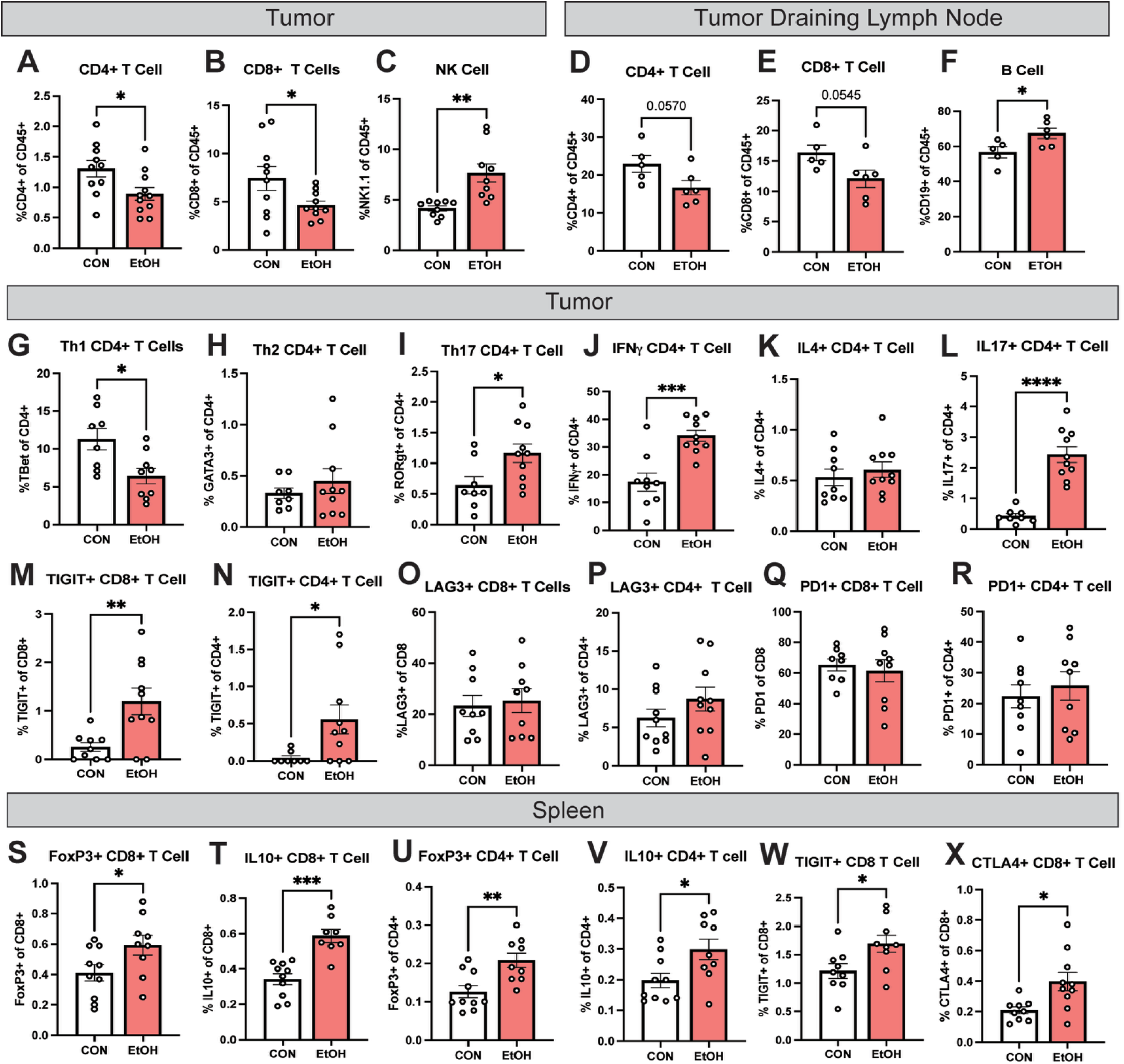
Ethanol disrupts Th1 CD4 differentiation and promotes regulatory T cell phenotypes. Mice were treated with EtOH (5g/kg) or water 5 days/week for 5 weeks followed by injection of MB49 (10^7^ cells, i.d.). At day 8 PTI, mice receive anti-PD1 (200 mg/kg, i.p.) with sacrifice at day 11 for flow cytometry of tumor, tumor draining lymph nodes, and spleen. In the tumor, **(A)** ethanol reduces the overall number of CD4+ T cells significantly and **(B)** modestly reduces the number of CD8+ T cells. **(C)** Ethanol increases the number of NK cells in the tumor. In the tumor draining lymph node there is an ethanol-induced reduction in both **(D)** CD4 and **(E)** CD8 T cells with an increase in **(F)** B cells. Ethanol reduced the number of **(G)** Th1 T cells had no change on **H)** Th2 and increased the proportion of **(I)** Th17 T cells with **(J)** measurable increases in IFNg. **(K)** No changes in the proportion of IL4+ CD4 T Cells. **(L)** There was a significant change in IL17+ CD4 T cells were found with an increase from less than 1% IL17+ to over 2%. Ethanol increased the proportion of **(M)** TIGIT+ CD8+ and **(N)** CD4+ T cells. Ethanol reduced the proportion of **(O)** LAG3+ CD8 T cells with no significant increases in **(P)** LAG3+ CD4, **(Q)** PD1+ CD8 or **(R)** PD1+ CD4 T cells. In the spleen ethanol increased the proportion of **(S)** FoxP3 CD4, **(T)** IL10+ CD4, **(U)** FoxP3 CD8, **(V)** IL10+ CD8, **(W)** CTLA4+ CD8, and **(X)** TIGIT+ CD8 T cells. All statistical testing was done with a two-sided unpaired t-test.

We next assessed specific adaptive immune cell phenotypes associated with ICI response. For CD4^+^ T cells, the Th1 is the preferred subtype for anti-tumor responses, while Th2 and Th17 are less productive and could in part promote a pro-tumor microenvironment *(36)*. Within the tumor, alcohol reduced the proportion of TBet+ Th1 CD4+ T cells (**Fig. 3G**), GATA3+ Th2 were unchanged (**Fig. 3H**), and RORγt+ Th17 CD4+ T cells were increased (**Fig. 3I**). Both IFNγ and the Th17-associated cytokine IL-17 were increased within CD4+ T cells, while IL4 (Th2) was unchanged **(Fig. 3J-L)**. Alcohol also altered immune checkpoint molecule expression on tumor-infiltrating lymphocytes, increasing TIGIT was on both CD8+ and CD4+ T cells **(Fig. 3M-N**), while LAG3 and PD1 were unchanged **(Fig. 3O-R**). In the spleen, ethanol promoted FoxP3+ regulatory CD4+ and CD8+ T cell phenotypes (**Fig. 3S-T**) with increased expression of the anti-inflammatory cytokine IL10 in both cell types (**Fig. 3U-V**). Similar to findings within the tumor there was increased checkpoint molecule expression suggesting an exhaustion-like phenotype in CD8+ T cells with increases in CTLA4 and TIGIT (**Fig. 3W-X)**. Thus, alcohol promotes a shift from anti-tumor Th1 to Th17 CD4+ T cell phenotypes, increases exhaustion-associated markers, and promotes regulatory phenotypes in the periphery.

### Alcohol disrupts CD8+ T cell activation and effector function in a cell autonomous manner

To define the underlying mechanisms of alcohol-induced T cell dysfunction, we profiled the transcriptomes of OT-1 CD8+ T cells with or without ethanol exposure in naïve and activated states. Alcohol altered the OT-1 transcriptome in the naïve unstimulated state, increasing 621 and decreasing 460 transcripts. Alcohol increased expression of T cell activation-associated genes including Egr1, Egr2, Fos, and FosB **(Fig. 4A)**. IPA predicted reductions in MYC signaling, increases in TP53 (**Fig. 4B**), and induction of T cell activation pathways IFNγ and NFAT **(Fig. 4C)**. In the context of TCR activation with CD3/CD28 beads, however, reduced T cell activation was seen with alcohol. Erg1, Fos, and FosB were reduced **(Fig. 4D)**, and IPA predicted suppression of IFNγ and IL-2 upstream signaling with increased MYC **(Fig. 4E)**, and reductions in T cell activation-associated pathways **(Fig. 4F)**. This indicates alcohol has cell autonomous effects on CD8+ T cell activation, the directionality of which depends on TCR stimulation. It also suggests alcohol promotes TCR-independent T cell activation that results in insufficient stimulation and/or exhaustion with subsequent TCR stimulation.

**Fig. 4:**
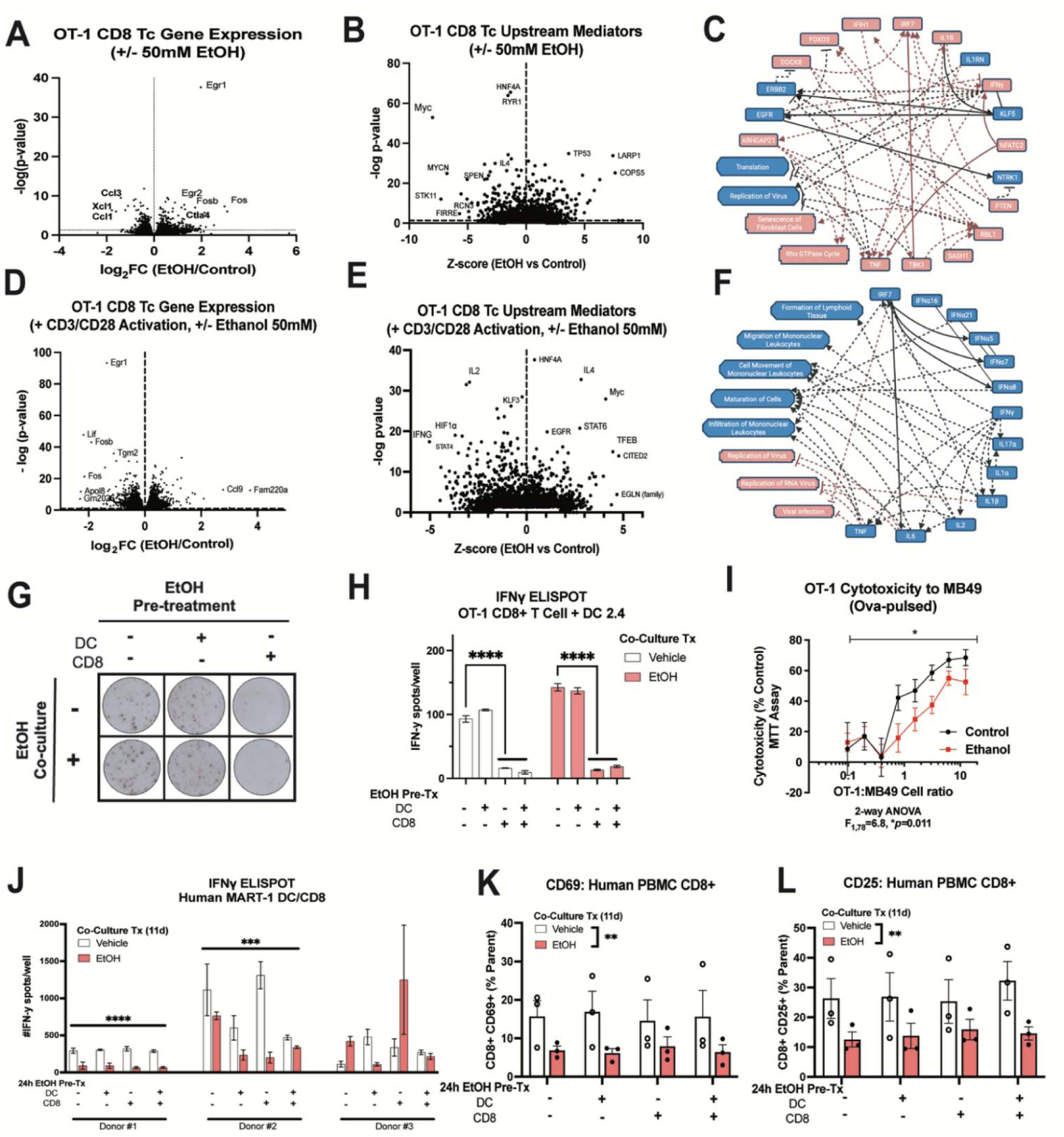
Pre-exposure of CD8 T cells to ethanol reduces activation. (A-F) OT-1 CD8+ T cells received ethanol (50mM) or vehicle for 96 hours +/-CD3/CD28 TCR activation followed by RNA-Seq (3 animals/group). **(A)** Ethanol increased T cell activation markers Egr1, Egr2, Fosb, and CTLA4. **(B)** IPA upstream mediator analysis predicted reduced MYC and increased TP53 signaling with ethanol. **(C)** Predicted functional impact of ethanol on OT-1 indicates increased activation markers IFN, TNF, IRF7, and IL1b. **(D)** Ethanol disrupted the CD3/CD28-induced transcriptional response reducing activation markers Egr1, Fosb, and Fos. **(E)** IPA upstream mediator analysis predicted increased MYC signaling and decreased IL2 and IFNg with ethanol in the setting of TCR activation. **(F)** IPA predicted reduced interferon and inflammatory signal in CD3/CD28 activated T cells that received ethanol. **(G-H)** ELISPOT of OT-1 CD8+ T cells with DC 2.4 antigen presenting cells shows no impact of 24-hour ethanol pre-treatment on DCs, but a significant decrease in IFNg when CD8+ T cells are pre-treated with ethanol. Co-culture of ethanol for 24 hours made no difference. (Anova, multiple comparisons test). **(I)** An MTT assay was done to measure OT-1 T cell killing efficiency when both T cells and tumor cells were exposed to ethanol (24 hour, 50mM) finding a significant (p=0.011) decrease in T cell killing upon ethanol exposure across several ratios of tumor and T cells (2-way ANOVA). **(J-L)** Human naïve CD8+ T cells and monocyte-derived dendritic cells were collected from 3 patients and pulsed with MART1. DC and T cells were either exposed to ethanol or not for 24 hours followed by a co-culture with ethanol exposure or not to allow for T cell maturation. **(J)** Two of the three donors had a significant decrease of IFN activity post 12-day co-culture with ethanol. Flow cytometry indicates a significant decrease in **(K)** CD69 and **(L)** CD25 (n= 3 patients, average of 3 technical replicates, 2-way ANOVA).

We next assessed the impact of alcohol on CD8+ T cell function. IFNγ secretion from OT-1 CD8+ T cells in response to cognate antigen presented by OVA-pulsed DC 2.4 cells was assessed by ELISPOT. DC 2.4 or OT-1 cells were pre-incubated +/-ethanol for 24 hours, followed by co-culture +/-ethanol. Preincubation of DCs with alcohol had no effect on T cell stimulation (**Fig. 4G**, top middle), neither did alcohol during co-culture without CD8+ T cell alcohol preincubation (**Fig. 4G**, bottom left and middle). However, a significant decrease in IFNγ production occurred when CD8+ T cells were preincubated with alcohol (**Fig. 4G**, right column and **Fig. 4H**). Ethanol also reduced OT-1 CD8+ T cell killing efficiency of OVA-pulse MB49 tumor cells (**Fig. 4I**, 2-way ANOVA, F_1,76_ = 6.8 *p* = 0.011). We then assessed functional responses of human CD8+ T cells from 3 HLA-A02+ donors stimulated with their cognate MART1 antigen. Naive CD8+ T cells and MART-1 pulsed DCs were co-cultured +/-ethanol during the 12-day T cell maturation period. In two of the three patients, alcohol reduced CD8+ T cell production of IFNγ **(Fig. 4J).** Flow cytometry found recent (CD69) and persistent (CD25) activation were decreased by ethanol **(Fig. 4K-L)**. Together, these findings indicate that alcohol alters CD8+ T cell function, reducing activation, IFNγ production, and T cell killing efficiency.

### Alcohol disrupts tissue resident T cell memory formation and promotes recurrence

Effective T cell memory formation is essential for durable responses to ICI. Therefore, we investigated the impact of alcohol on memory and recurrence in the MB49 model. Mice received 5 weeks of alcohol or water treatment followed by MB49 grafting and anti-PD1 ICI with assessment on post-tumor day 11 as in **Fig. 3**. In the tumor, alcohol did not significantly alter proportions of naïve (CD44+ CD62L-), effector (CD44-CD62L-), effector memory (CD44-CD62L+), or central memory (CD44+ CD62L+) CD8+ or CD4+ T cells (**Fig. 5A-B**). CD103, which binds E-cadherin to localize T cells peripheral tissue, had a trend toward a significant reduction with ethanol in CD8+ T cells with a significant reduction of CD103+ CD4+ T cells (**Fig. 5C-D**). Alcohol significantly reduced both CD8+ and CD4+ tissue resident T cells (CD44+ CD62L-CD69+) *(37, 38)* in the tumor (**Fig. 5E-F**). In the spleen, there were no significant changes to naïve, effector, effector memory, or central memory phenotypes in CD8+ or CD4+ T cells (**Fig. 5G-H**). However, similar to the tumor a reduction in CD103 was found in both CD4+ T cells and CD8+ T cells (**Fig. 5I-J**). Further, in both CD8+ and CD4+ T cells, we found a reduction in CCR7 surface expression, an important receptor for T cell trafficking to the lymph nodes and secondary lymphoid organs (**Fig. 5K-L**). Thus, alcohol disrupts the formation of tissue resident T cells without disrupting effector memory or central memory subsets in the context of MB49 bladder cancer with ICI.

**Fig. 5:**
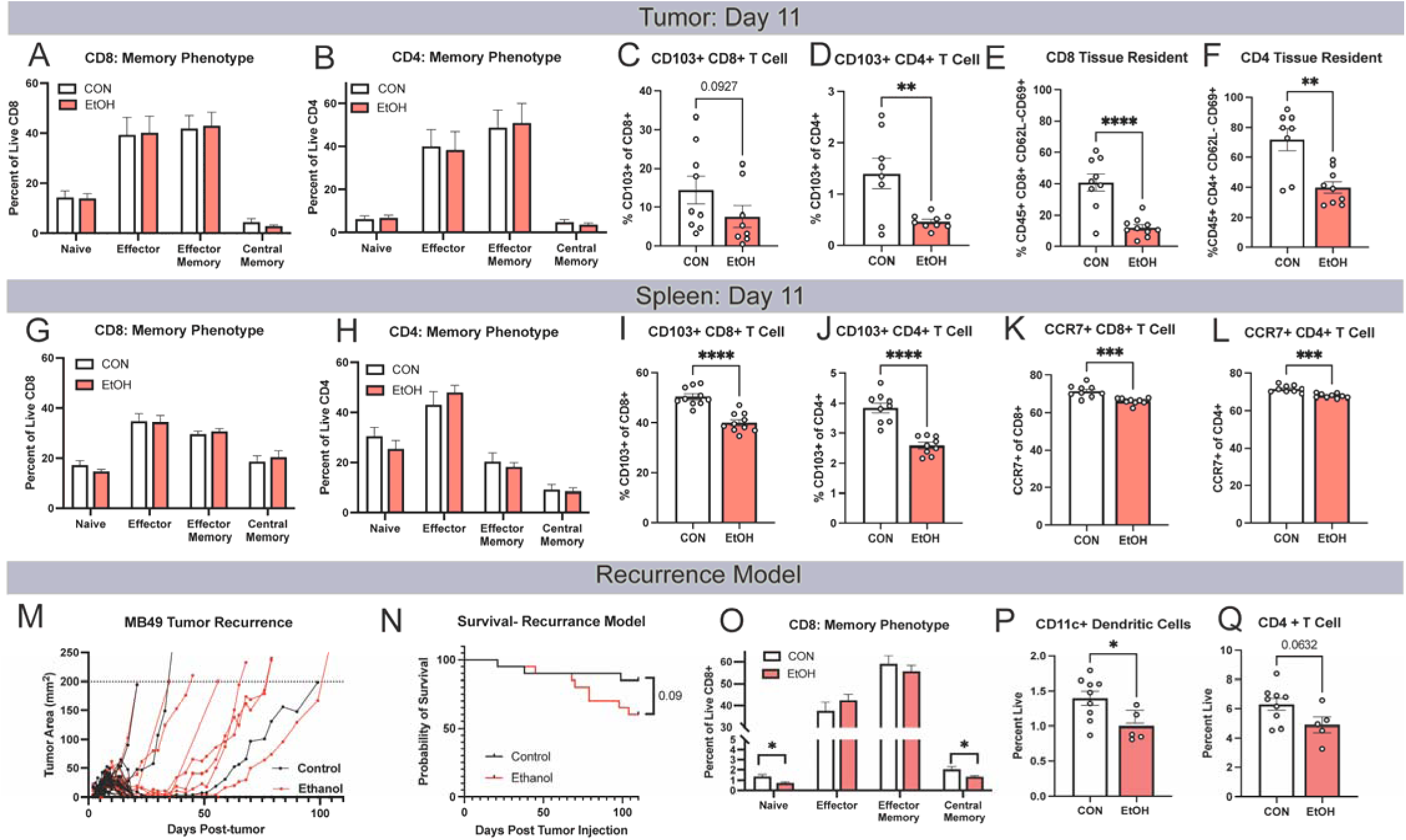
Ethanol reduces T cell memory formation resulting in increased relapse. (A-L) Mice were treated with EtOH (5g/kg) or water 5 days/week for 5 weeks followed by injection of MB49 (10^7^ cells, i.d.). At day 8 PTI, mice receive anti-PD1 (200 mg/kg, i.p.) with sacrifice at day 11 for flow cytometry of tumor, and spleen. There were no significant changes in the memory phenotypes of **(A)** CD8+ or **(B)** CD4+ memory in the tumor. **(C)** CD103 was significantly decreased in CD4+ T cells, and **(D)** non-significantly decreased in CD8+ T cells. **(E-F)** Tissue resident memory cells are also decreased in the ethanol treated mice in both CD4+ and CD8+. **(G-H)** In the spleen, we profiled CD8+ T cell phenotype and CD4+ T cell phenotype finding no significant differences. In CD103, we find a decrease in both **(I)** CD8+ and **(J)** CD4+. We also find decreases in CCR7 in both **(K)** CD8 and **(L)** CD4. **(M-Q)** We created a model to study recurrence in which C57Bl6/J mice were treated with EtOH or water control for 5 weeks (5 days/week, 5 g/kg i.g.) and then injected with a small “vaccine” dose of 10^6^ cells and anti-PD1 at day 8 PI. Tumor were monitored for moribund criteria and mice continued receiving ethanol daily. **(M)** Tumor burden was measured as mice began to recur **(N)** survival was non-significantly different between the ethanol and control group (p = 0.09, log-rank test). After the growth of tumors, mouse spleens were harvested from the remaining mice where we found a **(O)** significant decreases in CD8+ T naïve and central memory populations in ethanol treated mice with a non-significant increase in effector CD8+ T cells **(P)** a decrease in CD11c+ dendritic cells **(Q)** non-significant CD4+ T cell reduction (p-value = 0.632). All statistical tests unless otherwise indicated used an unpaired two-sided t-test.

In order to assess the impact of alcohol on functional immunological memory after initial response to ICI, we modified the MB49 ICI paradigm to promote complete response. Mice received 5 weeks of ethanol or water gavage pre-treatment followed by injection of a lower number of tumor cells (10^5^/mouse). Nearly all mice showed a complete response after anti-PD1 treatment. Alcohol increased the rate of spontaneous recurrence (**Fig. 5M**). All recurrences occurred by day 51 after tumor injection. After the 10-week monitoring period for recurrences, spleens were harvested from remaining mice and assessed by flow cytometry. With this prolonged ethanol exposure, a reduction of both naïve CD8+ T cells as well as CD8+ central memory T cells were found (**Fig. 5O**). Further, alcohol reduced the proportion of dendritic cells with a trend toward a reduction in splenic CD4+ T cells (**Fig. 5P-Q**). This indicates that alcohol reduces durable immunity and promotes recurrence after initial response to anti-PDI ICI in the MB49 bladder cancer model.

## Discussion

Alcohol is a commonly used and misused substance that has profound impacts on the immune system. Here we report that alcohol use reduces the efficacy of ICI immunotherapy. This was found in human patients, preclinical mouse models, and studies *ex-vivo* and *in vitro* using both mouse and human T cells. Patients with lung or bladder cancer who report regular alcohol use had mortality rates approximately twice those of patients who did not drink alcohol. Similar findings were observed *in vivo,* where alcohol reduced ICI efficacy by ∼50% in lung (LN4K1) and bladder (MB49) models. Alcohol disrupted the human and mouse tumor transcriptomes, tumoral T cell phenotypes, and T cell anti-tumor responses in manners predicted to reduce ICI efficacy. Together, these findings indicate that alcohol use is a detriment to ICI immunotherapy efficacy, warranting clinical recommendations for alcohol cessation during treatment, and further studies to identify approaches to reverse these effects.

Our assessment of outcomes of human patients found increased mortality for patients with lung or bladder cancer who regularly consumed alcohol. To our knowledge this is the first study to report a detrimental effect of alcohol use in human patients undergoing cancer treatment. In this retrospective study, we divided patients into two groups – occasional drinkers who reported less than 1 drink per week and regular drinkers who reported greater than 1 drink per week. There are some inherent limitations to this approach, though it likely reduces biases in alcohol self-reporting. Specifically, there is a known under-reporting bias for patients who consume alcohol, whereas patients who do not consume alcohol tend to be more accurate in their reporting *(39-41)*. This under-reporting bias makes it difficult to accurately quantify alcohol use retrospectively; limiting the accuracy of assessing the alcohol quantity consumed as a continuous variable. Further, since the study was retrospective, there was no way to validate the level of alcohol use biologically. Future prospective human clinical trials incorporating frequent alcohol intake questionnaires and biological assessment of recent alcohol use using biomarkers such as phosphatidyl ethanol are needed. However, even with this under-reporting bias, the level of drinking in regular drinkers (∼7-8 drinks/week) was at the level associated with risky/hazardous drinking defined by several regulatory bodies and the World Health Organization *(42, 43)*. It was interesting, and a bit confounding that no effect of alcohol was found in patients with head and neck cancer, the only cancer type in our dataset in which alcohol is the main carcinogen. This suggests possible inherent differences in the immune response for alcohol-induced cancers versus other cancer types, though this is yet unknown. Future prospective studies should more definitively establish the risk of alcohol use with different tumor types. It is also possible that other lifestyle behaviors from patients who abstain from alcohol could promote better outcomes. However, our findings in murine models of lung and bladder cancer strongly suggest that alcohol itself reduces ICI efficacy. It is unknown if regular alcohol use would also be detrimental for other forms of cancer therapy (e.g., chemotherapy or radiation). This should also be investigated in prospective clinical trials *(29)*.

Studies in mouse models of lung and bladder cancer were strikingly consistent with the retrospective human analysis. Interestingly, alcohol had no clear impact on tumor growth or survival in the absence of ICI. However, alcohol reduced survival in mice that received ICI both for lung and bladder cancer, implicating a detrimental effect of alcohol on anti-tumor immunity. This involved reductions in tumoral CD4+ and CD8+ T cells as well as a shift from Th1 to Th17 and less robustly Th2 phenotypes. The Th1 phenotype is generally considered necessary for effective anti-tumor immunity *(36, 44)*. For example, Th1 T cells promote the release of IFNγ which increases antigen presentation and inhibits angiogenesis *(45, 46)*. Th2 polarization is associated with pathological fibrosis and tumor progression in bladder *(47)*. In humans Th1 is associated with increased survival and Th17 with decreased survival in colorectal cancer *(48)*, while in pancreatic cancer, Th2 polarization is associated with reduced survival *(49)*. Th17 cytokines have been associated with immune related adverse events with anti-PD1 therapy *(50)*, and Th17 cells were reported to promote resistance to combined anti-PD-L1 and MEK inhibition *(51)*. In the spleen, we found an increase in regulatory CD4+ and CD8+ T cells, which could reduce ICI efficacy by releasing IL10, lysing effector T cells, or inhibiting antigen presenting cells *(52)*. Our findings are consistent with other studies that find alcohol favors the expression of Th2 cytokines in the setting of bacterial pneumonia *(53)* and work finding that chronic alcohol increases in both IFNγ and IL17 in the blood and spleen *(54)*. Further work is needed to elucidate to define the mechanism by which alcohol seems to promote Th2, Th17, and T regulatory phenotypes and to determine if alcohol exerts the same impact using genetically engineered mouse models.

Human and mouse *ex-vivo* as well as *in vitro* studies indicate that alcohol alters T cell phenotypes downstream of TCR signaling using both transcriptomics and functional assays. First, we observed that naïve T cells seem to have increased basal activation independent of the TCR when exposed to alcohol with RNAseq finding increased activation markers *(Fos*, *FosB*, *Egr1* and *Egr2)*. On the contrary, with TCR activation, these same genes were downregulated by alcohol. This could indicate a form of TCR-independent ‘bystander’ activation *(55)*by alcohol that results in an exhaustion-like phenotype with robust TCR stimulation. This is supported by findings *in vivo* where alcohol increased checkpoint inhibitors were in the tumor (TIGIT) and spleen (CTLA4) and may be underlying the increase in TCR signaling predicted in human lung tumors. It is currently unclear how alcohol is inducing its cell-autonomous effects on T cells. This might be through effects on activity-regulating kinases downstream of the TCR or through effects on T cell Toll Like Receptors (TLRs). Alcohol can autonomously activate kinases such as MAPK and PI3K in other cell types that regulate T cell activity as well as TLRs such as TLRs 2 and 4 *(56, 57)*. Further, it is possible that alcohol alters TCR activation by changing membrane fluidity. A prior study found that ethanol disrupts the TCR stimulation with decreased localization of Lck and LAT to the membrane *(58)*. Future studies will investigate the molecular and membrane effects of alcohol on T cells, and how this determines the T cell phenotypes in the setting of neoantigen stimulation.

In addition to detrimental impacts on anti-tumor immunity, alcohol also had effects on human and bladder tumor cells. In both human tumor bulk RNAseq and murine MB49 bladder cells, alcohol altered the transcriptional responses, yielding a decrease in pro-inflammatory and immune-related pathways such as IFN signaling that activate anti-tumor immunity. Consistent with this idea, a previous study found that tumors that have lost the IFNγ signaling are more likely to be resistant to anti-CTLA4 ICI immunotherapy *(59)*. Further, IPA predicted increases in oxidative phosphorylation (OXPHOS) in both human lung tumors and murine MB49 cells treated with alcohol. Recently, tumor OXPHOS has been inversely linked with antigen presentation, with inhibition of tumoral OXPHOS increasing tumor antigen presentation and immunogenicity*(35)*. Previous studies have shown that tumors relying on oxidative phosphorylation have an increased risk of metastasis and are more resistant to treatment *(60, 61)*. Further, in recent studies tumor OXPHOS has been inversely linked with antigen presentation, with inhibition of tumoral OXPHOS increasing tumor antigen presentation, immunoproteasome subunit expression, and immunogenicity *(34, 35)*. This inverse relationship between tumor OXPHOS and antigen presentation could underlie our observed reduction in MHC machinery and key immunoproteasome components in MB49. The impact of alcohol on mitochondria and OXPHOS can vary depending on the cell type and context *(62, 63)*. Alcohol is metabolized to acetate, which upon conversion to acetyl-CoA can enter the citric acid cycle to promote OXPHOS. However, the impact of alcohol on tumor metabolism across different tumor subtypes is understudied and a focus on ongoing investigation. These studies will determine if alcohol reduces tumor immunogenicity through loss of interferon signaling and induction of tumoral OXPHOS.

We also found that alcohol disrupts aspects of immunological memory. Alcohol exposure reduced the expression of tumor and splenic CD103 and splenic CCR7, which promote tissue resident memory and lymphoid recruitment during re-exposure respectively *(64, 65)*. This finding is consistent with a previous study that reported an ethanol-induced decrease in CD103 after 6-weeks of alcohol intake *(66)*. A recent meta-analysis revealed that CD103+ immune cells in the tumor is associated with increased overall survival, and disease-free survival *(67)*. Thus, the loss of CD103 with alcohol could contribute to the increased recurrence we observed in our MB49 recurrence model. A similar version of our recurrence model found durable memory is heavily reliant upon CD4+ T cells *(68)*. We also found a reduction in central memory CD4+ T cells in our model which could also contribute to recurrence. Previous studies have shown that central memory may be even more effective than effector memory T cells due to a heightened recall response and a prolonged lifespan *(69, 70)*. Given these findings, future human studies should also monitor the rate of recurrence as well as the formation of memory T cell subtype in patients who regularly consume alcohol.

According to the Cancer Research Institute Impact Report 2024, about 9 million cancer patients (45% of new cancers) in the United States are eligible for immunotherapy. This equates to nearly 7 million people that may receive immunotherapy that currently drink alcohol, with about 3.5 million people drinking at hazardous levels. Our work suggests that the outcomes of up to 3.5 million people could be improved by them ending their alcohol use. It also suggests that recommendations against drinking and interventions that reduce drinking in this high stress population could improve survival. As stated above, more studies will be needed to define the exact mechanism underlying alcohol-induced T cell dysfunction and loss of ICI response, to firmly establish the dose-response relationship, and to identify therapeutic interventions for patients with difficulty stopping alcohol use. Future studies should also assess the effect of alcohol on other modalities of immunotherapy such as chimeric antigen responsive T cells (CART), tumor antigen vaccines, radiation therapy, and chemotherapy. Nonetheless, this current work implicates alcohol as a negative determinant of ICI immunotherapy efficacy that should be further investigated and potentially communicated to patients.

## Methods

### Human Survival and Alcohol Use Data Provenance

Patient demographics, survival, and alcohol use data from a previous study of patients receiving the anti-PD1 drugs nivolumab or pembrolizumab were obtained based on a protocol approved by the UNC Institutional Review Board(#LCCC1520). Patients were grouped into binary categories of nondrinkers (≤1 drink/week) or drinkers (>1 drink/week) based on alcohol consumption recorded in the medical chart.

### Mouse Models

The University of North Carolina (UNC) Institutional Animal Care and Use Committee approved the protocol for this experiment (IACUC #23-099 and #23-230).

The experimental environment consisted of a temperature and moisture regulated room that maintained a 12-hour light/dark cycle. We obtained female C57Bl/6 mice from Charles River Laboratory and allowed them to habituate for 1 week. Upon receipt, we placed 5 mice per cage and allowed ad libitum access to food and water for the duration of the experimental period. For the pre-treatment period (5 weeks), we administered 2.5g/kg ethanol, 5.0g/kg ethanol, or water vehicle by oral gavage to mice. We continued oral gavage for the remainder of the study.

After the pretreatment period, mice received orthotopic injections to the intrapulmonary space as previously reported*(32)*. Briefly, mice were placed under anesthesia and an incision was made between rib 10 and 11 and 10^5^ cells (in HBSS) were infused into the right lung parenchyma. Mice received anti-PD1 antibodies at day 8,11, and 14 (200 ug/ml) and anti-CTLA4 (200 ug/ml) every 3 days throughout the experiment. We monitored the mice until morbidity criteria were reached (tumor area > 200 mm^2^, > 10% body weight loss) and then humanely euthanized.

MB49 bladder tumors were treated similarly in which after the pre-treatment period, we injected mice subcutaneously with 10^7^ MB49 murine bladder cancer cells suspended in PBS. Tumor size was collected every 2-3 days and murine body weight was collected weekly. On days 8,11, and 14 after tumor instillation, we treated mice with 200 ug/ml anti-PD1 antibodies (clone: J43, BioXcell, Hanover, NH) in 100ul sterile phosphate-buffered saline (PBS). We monitored the mice until morbidity criteria were reached (tumor area > 200 mm^2^, > 10% body weight loss) and then humanely euthanized. Multiple cohorts were assessed which were assessed for consistency across experiments in a meta-analysis adjacent forest plot (**Fig. S3A-B**). For the recurrence model, pre-treatment remained the same, but mice received 10^6^ MB49 tumor cells and 1 day treatment (day 8) of anti-PD1 antibodies (200 ug/ml).

### Flow Cytometry

We treated mice for flow cytometry as described above, except that instead of receiving 3 doses of antibodies, they received only one dose on day 8. On day 11, we sacrificed the mice and collected tumor, spleen, and tumor-draining lymph nodes. Tissues were processed into single-cell suspensions using the MacsQuant C-tube and dissociator (Miltenyi Biotec, Auburn, CA) and filtered with a 70um filter. Red blood cells were removed using 1mL of ammonium-chloride-potassium (ACK) lysis buffer (Thermo Fisher Scientific, Waltham, MA) for two minutes followed by quenching with 9mL of RPMI media. Suspensions were then centrifuged at 1200rpm for 5 minutes and filtered through 40µM filters. Single cell suspensions were counted on the Countess II (Fisher) and plated in a 96-well staining plates at about 2 million cells per well. Cells were stained for viability using Fixable Viability Stain 780 (1:1000 in PBS) for 20 minutes at 4C. Cells were then washed with staining buffer (RnD Systems) followed by FC receptor block (1:100 in staining buffer) for 15 minutes prior to addition of extracellular antibodies for 1 hour. Cells were then fixed and permeabilized for 10 minutes (RnD Systems) and washed twice with permalization/wash buffer and resuspended into intracellular antibodies overnight.

Samples were run on either the BD Symphony for conventional flow or the Cytek Aurora for spectral flow. Analysis for percent of positive cells was completed using FlowJo (Ashland, OR). Dead cells and doublets were removed prior to analysis.

### Cell culture

After processing OT-1 T cells as previously described, we isolated CD8+ T cells using a negative selection magnetic bead-based kit (Stem Cell Technologies, Cambridge, MA). We next cultured the isolated T cells in complete Roswell Park Memorial Institute (RPMI) media supplemented with 10% fetal bovine serum (FBS, Corning), 1% penicillin-streptavidin (Gibco), and 1x β-mercaptoethanol. For every 2 days in culture, we added 100 units/mL of IL-2 to each culture plate promote cell growth and prevent apoptosis. To activate OT-1 T cells, we utilized CD3 (1 ug/ml) and CD28 antibodies (1 ug/ml) diluted in PBS and allowed to incubate in a 24-well non-tissue culture treated plate overnight. We then removed the PBS and applied cells for the duration of the activation period.

MB49 murine cell line tumors which we cultured in Dulbecco’s Modified Eagle Medium (DMEM) supplemented with 10% FBS (Corning) and 1% penicillin-streptavidin (Gibco). Cells were split 1:5 every 2-3 days.

### MTT Assay

MB49 bladder tumor cells were plated at 10,000 cells/well in a 96 well plate after being exposed to ethanol (50mM) for 24 hours. The MB49 cells were pulsed with SIINFEKL (20 ug/ml). OT-1 CD8+ T Cells were also exposed to 50mM ethanol for 24 hours. OT-1 T cells are plated using a 96 well plate and completing serial dilutions from 25 T cells: 1 tumor down to 1 T cell: 5 tumor cells. For controls, a max kill well used triton X to kill all tumor cells and a No kill well had only tumors and no T cells. T cells and tumor cells were allowed to interact overnight and then the MTT assay (Abcam) was run according to standard protocol.

### ELISPOT

#### Human

We ordered eight leukopaks from Gulf Coast Regional Blood Center and completed HLA typing by flow cytometry for HLA-A02+ samples, we had 3 HLA-A02+ samples. For HLA-A02+ samples, we performed PBMC isolation by Ficoll density-gradient centrifugation. We utilized approximately 1/3 of the PBMCs to start the DC cultures using plastic adherence property of monocytes. DC cultures were started in two different groups for each blood sample, one control group and one ethanol treated group (50mM ethanol). The alcohol treated DC culture flasks were kept in a consistent state by remaining in an incubator with 2% ethanol in the environment. The remaining 2/3 of PBMCs were cryopreserved for naive CD8+ isolation. Monocytes were differentiated into immature DCs in complete RPMI media containing 10% human serum (Millipore, H4522), penicillin streptomycin (Gibco, 15070063) and L-glutamine (Gibco, 25030081), in the presence of GM-CSF (1000 IU/mL, RnD Systems) and IL-4 (1000 IU/mL, RnD Systems) during a 6-day culture and then into mature DCs in presence of IFN-y (100 IU/mL, RnD Systems) and LPS (10ng/mL) overnight.

Frozen PBMC cells were thawed and used to isolate naive CD8+ T cells using a Stem Cell Technologies Naïve CD8+ T cell isolation kit (StemCell, 19258), they were then divided into two flasks for overnight culture with IL-7 and ethanol (50mM). We then begin co-cultures with the matured/pulsed DCs and nCD8+s at ratio of 4:1 (200K nCD8+:50K DC) in 200uL complete RPMI media, in presence of IL-21 (30 ng/mL), in round-bottom 96well plates. The media was replaced every other day (day 3, 5, 7, 9) by gently removing half the media from each well and adding fresh warmed media containing IL-7 & IL-15 (both 10 ng/mL, which makes final concentration of 5 ng/ml in culture well). The alcohol treated co-culture plate was kept in a consistent state by remaining in an incubator with 2% ethanol in the environment. While replacing the media for this plate, ethanol was added to the complete culture media with concentration of 5mM. After 12 days of co-culture, we performed an ELISPOT according to manufactures instructions utilizing PHA (Thermo Scientific, R30852801) as a positive control.

#### Murine

We used pooled OT-I mouse spleens and extracted CD8+T cells using a Stem Cell Technologies Kit. For antigen presenting cells, we utilized the DC2.4 cells will be used. CD8+ T cells and DC 2.4s were incubated +/-50mM ethanol for 24 hours. We then did a co-culture with or without 50mM ethanol for an additional 24 hours. Then the DC2.4 cells will be pulsed with OVA257-264 (SIINFEKL) peptide with final concentration of 10 mM/m. The ELISPOT was then completed according to manufactures instructions and read on the AID Classic ELR07 (Advanced Imaging Devices).

#### RNA Sequencing

We performed RNA sequencing on human tumor samples, OT-1 murine T cells, and MB49 murine bladder cell line. OT-1 T cells were exposed with or without CD3/CD28 activating antibody beads (DynaBeads) for 4 days with or without ethanol (50mM) exposure. MB49 cell lines were exposed with or without ethanol (25mM and 50mM) for 24 hours prior to library preparation. Human samples were paraffin embedded, and library preparations were done with Illumina FFPE library preparation kit.

Library preparations were prepared by the Immune Monitoring and Genomics Facility of the Lineberger Cancer Center. Sequencing was performed on short read Illumina technologies. Fastq files were aligned to the GRCh38 and GRCm39 for human and mouse, respectfully using STAR. Followed by quantification of RNA transcript abundance by Salmon. In MB49 and OT-1 T cells differential gene expression was found utilizing R package DESeq2. For human, since there was greater variation between the samples they were normalized by quantile normalization and fit to a linear model using R package limma. Ingenuity Pathway Analysis (IPA, Qiagen, Germantown, MA) was used to understand upstream mediators of RNA expression and graphical summaries of transcriptomic changes.

#### Statistical Analysis

We utilized Kaplan Meier survival curves for both human and murine survival. Cox regression of each individual tumor site was also completed with age, sex, and BMI used as covariates. Smoking was an additional covariate with results shown with and without smoking. Unless otherwise indicated, a two-sided t-test was performed in GraphPad Prism 10 (Boston, MA). For experiments utilizing both T cells and DCs an ANVOA was run.

## Acknowledgements

We would like to thank the Bowles Center for Alcohol Studies and the Flow Cytometry Core Facility (RRID: *SCR_019170)* for their assistance.

## Author Contributions

Conceptualization: LGC, BGV Data Curation: AK, KNG

Methodology: CP, LGC, BGV Investigation: KNG, JG, WB, MS, BC Visualization: KNG, LGC, BGV Funding acquisition: LGC, BGV Supervision: LGC, BGV

Writing – original draft: KNG, LGC

Writing – review & editing: KNG, LGC, BGV

