## Supplemental for "Alcohol use reduces the efficacy of anti-PD1 immunotherapy by disrupting anti-tumor immunity"

### Supplemental Data

| **Table S1: Human Subject Demographics** | | | |
| --- | --- | --- | --- |
| **Cancer Type** | **Occasional Drinkers** | **Regular Drinkers** | **P-value** |
|  | **Mean (SD)** | **Mean (SD)** |  |
| Lung (N = 92) |  |  |  |
| Age | 65.45 (11.28) | 64.03 (10.25) | 0.531 |
| Sex (% male) | 45.76% | 50% | 0.683 |
| Drinks/Week | 0.083 (0.279) | 7.367 (6.52) | **<0.001** |
| BMI | 24.27 (6.58) | 23.71 (5.90) | 0.705 |
| Smoking (%) | 91.07 | 100 | 0.059 |
| Bladder (N = 40) |  |  |  |
| Age | 69.19 (9.43) | 72.77 (7.66) | 0.244 |
| Sex (% male) | 21.43% | 53.84% | **0.048** |
| Drinks/Week | 0.17 (0.38) | 8.58 (6.63) | **0.0011** |
| BMI | 22.55 (5.27) | 19.97 (3.10) | 0.133 |
| Smoking (%) | 69.23 | 69.23 | 1 |
| Melanoma (N = 63) |  |  |  |
| Age | 62.97 (14.61) | 60.89 (15.19) | 0.497 |
| Sex | 64.29% | 60.71% | 0.762 |
| Drinks/Week | 0.24 (0.43) | 6.29 (4.59) | **<0.001** |
| BMI | 28.30 (6.01) | 27.94 (6.88) | 0.936 |
| Smoking (%) | 62.86 | 44 | 0.316 |
| Head & Neck (N = 50) |  |  |  |
| Age | 64.56 (10.21) | 60.5 (10.09) | 0.143 |
| Sex | 74.07% | 83.33% | 0.422 |
| Drinks/Week | 0.08 (0.28) | 9.13 (6.41) | **<0.001** |
| BMI | 24.47 (7.75) | 23.31 (7.85) | 0.712 |
| Smoking (%) | 62.93% | 92% | **0.013** |

| **Table S2. Impact of Alcohol Status on Overall Survival from Diagnosis.** Multivariable Regression Analysis. | | | |
| --- | --- | --- | --- |
| **Cancer type** | **Co-variate** | **β coefficient** | ***p*-value** |
| Lung  (N=93) | Age  Sex  BMI  Smoking Status  **Alcohol Use** | 0.018  -0.013  0.027  0.57  **-0.73** | 0.1448  0.9569  0.2153  0.2793  **0.0037**** |
| Bladder  (N=36) | Age  Sex  BMI  Smoking Status  **Alcohol Use** | 0.01  -0.15  0.026  0.40  **-0.55** | 0.42  0.63  0.25  0.08  **0.04*** |
| Melanoma  (N=67) | Age  Sex  BMI  Smoking Status  Alcohol Use | 0.0087  0.21  -0.02  0.019  -0.31 | 0.18  0.29  0.19  0.93  0.09 |
| Head and Neck  (N=50) | Age  Sex  BMI  Smoking Status  Alcohol Use | -0.0032  -0.077  0.013  -0.036  0.26 | 0.82  0.81  0.57  0.91  0.70 |

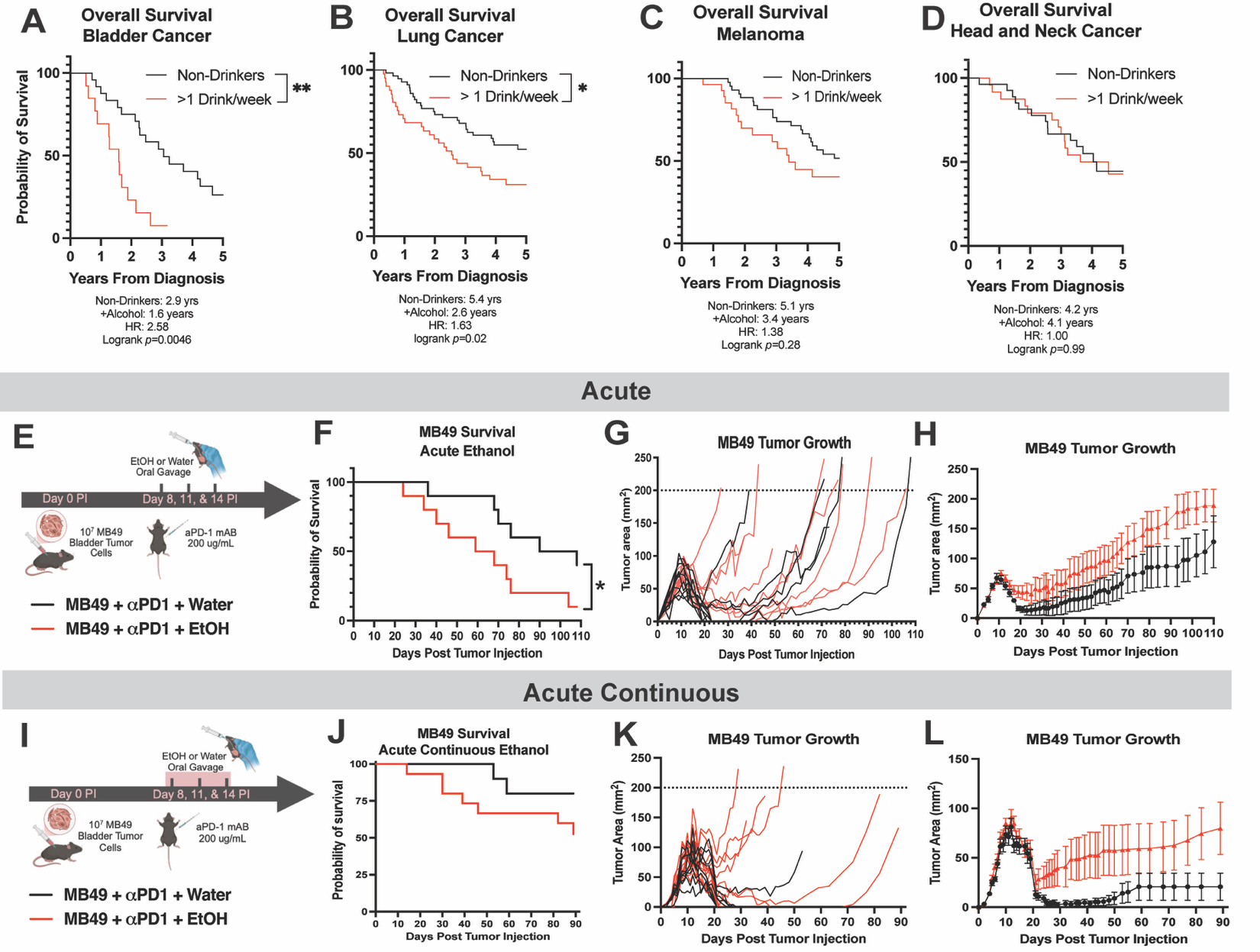

**Supplemental Fig. 1. Additional Human Survival Analysis and Murine Models**. **(A-D)** Retrospective analysis of patients treated with ICI who were nondrinkers (≤1 drink/week) or drinkers (>1 drink/week). **(A)** Patients with bladder cancer who drank alcohol had worse survival from diagnosis than non-drinkers (HR:2.58, Log-rank test). **(B)** Patients with lung cancer who drank alcohol had worse survival from diagnosis than non-drinkers (HR: 2.27, Log-rank test). **(C)** Patients with melanoma who drank alcohol did not have significantly worse survival from diagnosis than non-drinkers (HR: 1.38, Log-rank test). **(D)** There was no difference in survival between patients who drank or did not in head and neck cancer (HR: 1.00, Log-rank test). **(E)** The acute model received 10^7^ MB49 tumor cells and anti-PD1 ICI and EtOH (5g/kg, i.g. or vehicle) on day 8,11, and 14. **(F)** Acute alcohol treatment decreased survival to MB49 tumors. **(G-H)** Individual and average MB49 tumor burden. **(I)** The acute continuous model received 10^7^ MB49 tumor cells and anti-PD1 ICI on day 8,11, and 14, EtOH (5g/kg or vehicle) given day 8-14. **(J)** Acute continuous alcohol treatment had a non-significant decrease to survival to MB49 tumors (p = 0.1602, Log-rank test). **(K-L)** Individual and average MB49 tumor burden.

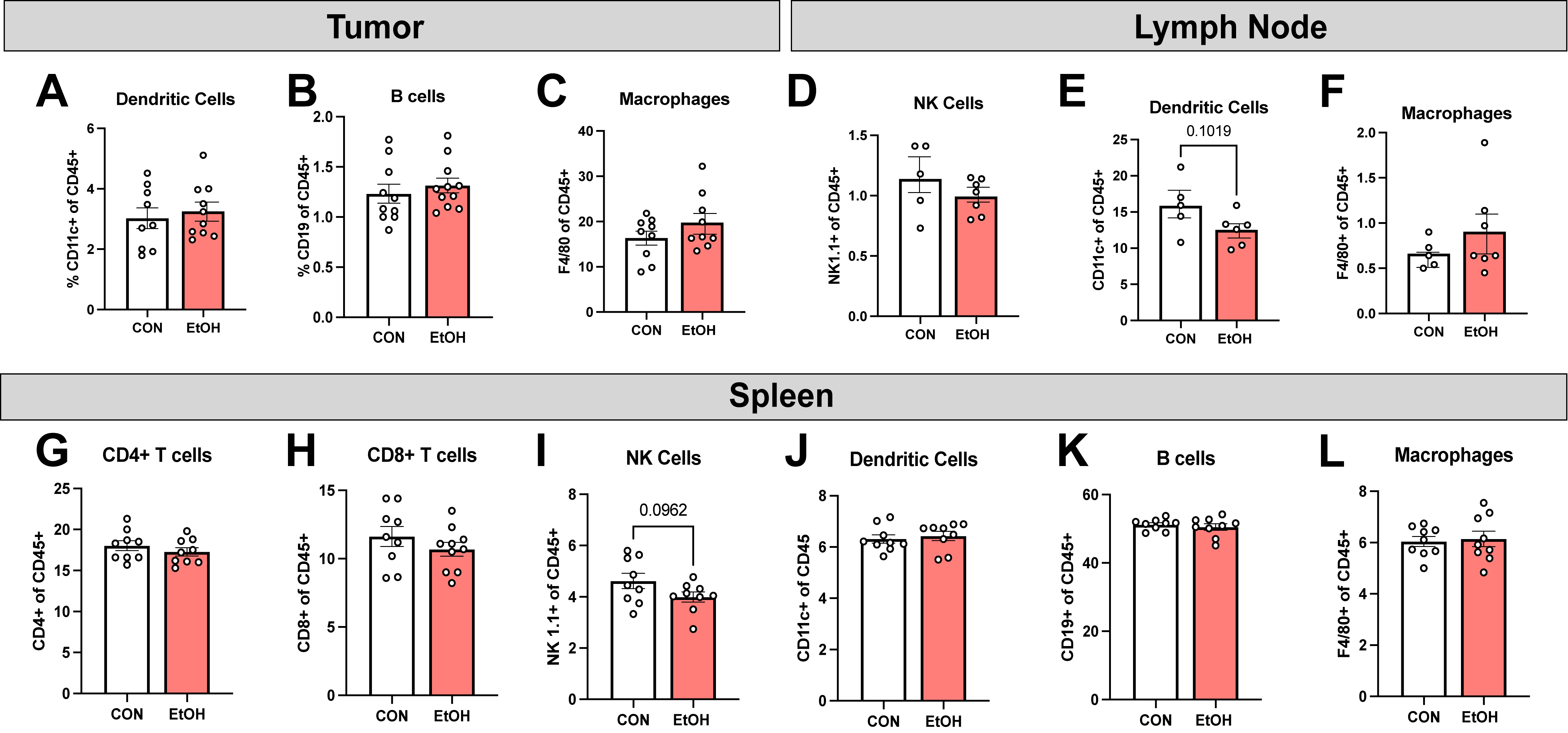

**Supplemental Fig. 2. Tumor, Tumor Draining Lymph Node, and Spleen Phenotyping.** (**A-L)** Mice were treated with EtOH (5g/kg) or water 5 days/week for 5 weeks followed by injection of MB49 (10^7^ cells, i.d.). At day 8 PTI, mice receive anti-PD1 (200 mg/kg, i.p.) with sacrifice at day 11 for flow cytometry of tumor, tumor draining lymph node, and spleen. Tumor has no change in **(A)** dendritic cells, **(B)** B cells, or **(C)** macrophages (two-tailed unpaired T test). Tumor draining lymph node had no changes in **(D)** NK cells, **(E)** dendritic cells, or **(F)** macrophages. Spleen had no changes in **(G-H)** CD4+ or CD8+ T cells, **(I)** NK cells, **(J)** dendritic cells, **(K)** B cells, or **(L)** macrophages.

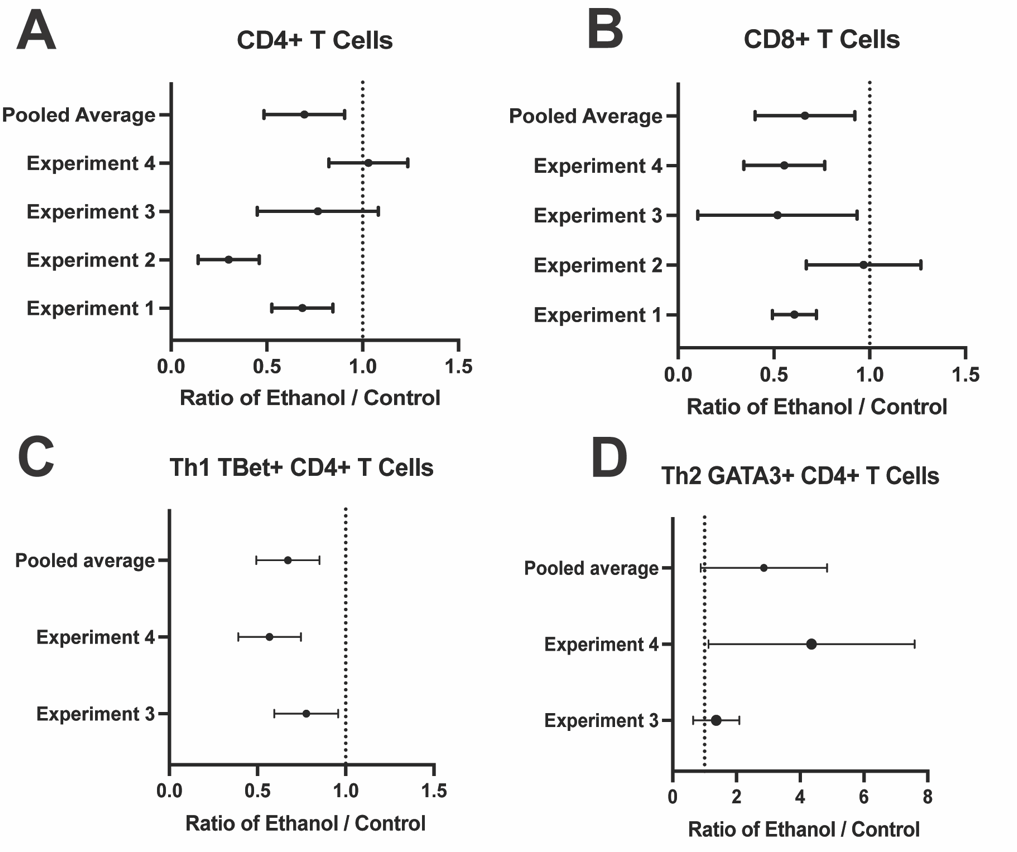

**Supplemental Fig. 3. Forest Plots of Multiple Flow Experiments. (A-D)** Mice were treated with EtOH (5g/kg) or water 5 days/week for 5 weeks followed by injection of MB49 (10^7^ cells, i.d.). At day 8 PTI, mice receive anti-PD1 (200 mg/kg, i.p.) with sacrifice at day 11 for flow cytometry of tumor. These experiments were completed independently up to 4 times and the odds ratio of ethanol / control for each experiment is graphed for **(A)** CD4+ T cells, **(B)** CD8 + T cells, and **(C)** CD4+ TBet+ T cells, and **(D)** CD4+ GATA3+ T cells.
